# Reprogramming and redifferentiation of mucosal-associated invariant T cells reveals tumor inhibitory activity

**DOI:** 10.1101/2021.06.21.449226

**Authors:** Chie Sugimoto, Yukie Murakami, Eisuke Ishi, Hiroyoshi Fujita, Hiroshi Wakao

**Author notes:** Corresponding author C.S.

## Abstract

Mucosal-associated invariant T (MAIT) cells belong to a family of innate-like T cells that bridge innate and adaptive immunities. Although MAIT cells have been implicated in tumor immunity, it currently remains unclear whether they function as tumor promoting or inhibitory cells. Therefore, we herein used induced pluripotent cell (iPSC) technology to investigate this issue. Murine MAIT cells were reprogrammed into iPSCs and redifferentiated towards MAIT-like cells (m-reMAIT cells). m-reMAIT cells were activated by an agonist and MR1-tetramer, a reagent to detect MAIT cells, in the presence and absence of antigen-presenting cells. This activation accompanied protein tyrosine phosphorylation and the production of T helper (Th)1-, Th2-, and Th17-cytokines and inflammatory chemokines. Upon adoptive transfer, m-reMAIT cells migrated to different organs with maturation in mice. Furthermore, m-reMAIT cells prolonged mouse survival upon tumor inoculation through the NK cell-mediated reinforcement of cytolytic activity. Collectively, the present results demonstrated the utility and role of m-reMAIT cells in tumor immunity, and will contribute to insights into the function of MAIT cells in immunity.

## Introduction

Mucosal-associated invariant T (MAIT) cells belong to a family of innate-like T cells harboring semi-invariant T-cell receptors (TCR), and recognize vitamin B2 metabolites as antigens on major histocompatibility complex class I-related gene protein (MR1) (Godfrey et al, 2019). MAIT cells play a pivotal role in immunity by bridging innate and adaptive immunities. They are abundant in humans, but rare in mice, and are associated with a number of diseases, such as bacterial and viral infections, autoimmune, inflammatory, and metabolic diseases, asthma, and cancer (Godfrey et al, 2015, Godfrey et al, 2019, Toubal et al, 2019). Although previous studies implicated MAIT cells in various tumors, their role in tumor immunity remains obscure. An analysis of infiltrating immune cells revealed that the high infiltration of CD8^+^CD161^+^T cells, largely comprising MAIT cells, in tumors represented a favorable prognostic signature across a wide array of human cancers (Gentles et al, 2015). In mucosal-associated cancers, including lung and colon cancers, increases in MAIT cells in tumors were concomitant with a decrease in circulation (Ling et al, 2016, Sundstrom et al, 2015, Won et al, 2016). While the high infiltration of MAIT cells in tumors negatively correlated with a favorable outcome in colorectal cancer (CRC) patients, the low infiltration of these cells in the tumors of hepatocellular carcinoma (HCC) patients correlated with a poor prognosis, which is in contrast to the role of MAIT cells in tumor immunity (Zabijak et al, 2015, Zheng et al, 2017).

Based on these discrepancies, further studies are warranted to investigate the role of MAIT cells in tumor immunity in mice. However, difficulties are still associated with examining MAIT cells in mice for the following reasons. These cells are 10- to 100-fold less abundant in mice than in humans. Although the advent of MR1-tetramer (MR1-tet) has enabled the detection of murine MAIT cells, the paucity of these cells *per se* still hampers functional studies (Corbett et al, 2014, Rahimpour et al, 2015). Furthermore, there is a lack of appropriate mouse models to investigate the function of MAIT cells (Garner et al, 2018, Godfrey et al, 2019, Toubal et al, 2019). Although the development of MAIT cells is dependent on MR1, MR1-deficient mice do not necessarily reveal all aspects of MAIT cells because they are also devoid of other MR1-dependent cells and provide only limited information on immune cells that potentially interact with MAIT cells. Similarly, previous studies on MAIT cell-specific TCR transgenic mice revealed the protective roles of MAIT cells in bacterial infections, type I diabetes, and experimental autoimmune encephalitis (Chua et al, 2011, Croxford et al, 2006, Cui et al, 2015, Le Bourhis et al, 2010, Martin et al, 2009, Meierovics et al, 2013, Reantragoon et al, 2013, Sakala et al, 2015, Shalapour et al, 2012, Shimamura et al, 2011). However, the aberrant expression of transcription factors in MAIT cells impedes investigations on their precise roles in health and disease.

We herein used induced pluripotent cell (iPSC) technology to examine the role(s) of MAIT cells in immunity, particularly tumor immunity. We reprogrammed murine MAIT cells into iPSCs (referred to as MAIT-iPSCs), differentiated MAIT-iPSCs into MAIT-like cells (referred to as m-reMAIT cells), and adoptively transferred them into syngeneic immunocompetent mice to investigate their role. This approach not only allows us to assess the role of m-reMAIT cells as effector cells, but also provides insights into interactions with other immune cells in tumor immunity.

In the present study, we showed that 5-(2-oxopropylideneamino)-6-D-ribitylaminouracil (5-OP-RU), an agonist of MAIT cells, and 5-OP-RU-loaded murine MR1-tetramer (mMR1-tet) both activated m-reMAIT cells, while these reagents induced the production of a similar, but not identical, set of cytokines and chemokines in the absence of antigen-presenting cells (APCs). When adoptively transferred, m-reMAIT cells migrated to different organs concomitant with the acquisition of maturation and improved mouse survival upon tumor inoculation. This tumor inhibitory activity was mediated through enhancements in the cytolytic activity of m-reMAIT cells by NK cells.

The present results imply that the adoptive transfer of m-reMAIT cells has potential as a novel tool for examining the immune functions of MAIT cells in health and disease.

## Results

### MAIT cell reprogramming, redifferentiation, and characterization of m-reMAIT cells

Fluorescent-activated cell sorting (FACS)-purified MAIT cells (defined as CD44^+^TCRβ^+^ mMR1-tet^+^ cells, purity >97%) from the lungs of C57BL/6 mice were reprogrammed with the Sendai virus vector KOSM302L (Wakao et al, 2013). Forty-six iPSCs were obtained with a reprogramming efficiency of 0.6%. Specific iPSCs were subjected to limiting dilutions and clone L7-1 was used throughout experiments unless otherwise indicated. Since TCR comprises α and β chains, the presence of rearranged *TRAV1-TRAJ33* specific for MAIT cells was detected by PCR followed by DNA sequencing and a Southern blot analysis with MAIT cell-derived iPSCs (MAIT-iPSCs) (Figures S1A-D). MAIT-iPSCs also harbored rearranged *TRBV13-3*, *TRBV19*, and *TRBV12-1* (Table 1) and exhibited pluripotency, as evidenced by the ability to give rise to chimeric mice (Figure S1E).

**Table 1.**
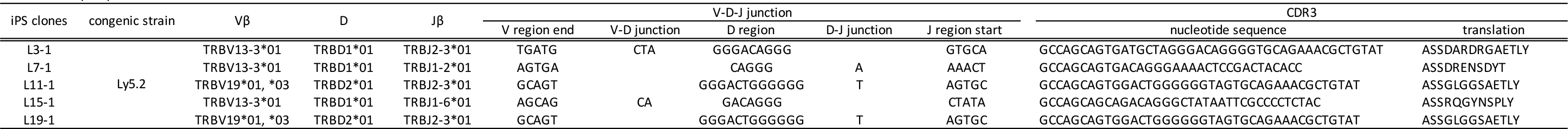
TRVB repertoires of MAIT-iPSCs. V, D, J, V-D-J junction, and CDR3 sequences of L3-1, L7-1, L11-1, L15-1, and L19-1 are summarized.

The differentiation of MAIT-iPSCs on OP9 ectopically expressing delta-like 1 (OP9-DLL1), a feeder cell for T-cell differentiation, resulted in m-reMAIT cells that were TCRβ^+^mMR1-tet^+^CD8^+^CD4^+^CD44^-^CD25^int^PLZF^-^RORγt^+^(Figure 1A). Consistent with this result, other MAIT-iPSC clones, such as L3-1, L11-1, L15-1, and L19-1, showed similar expression profiles to m-reMAIT cells on day 23 (Figure S1H). m-reMAIT cells multiplied more than 100-fold starting from MAIT-iPSCs with a doubling time ∼10.5 h during the logarithmic phase of growth later in differentiation (Figures S1F-G).

**Figure 1.**
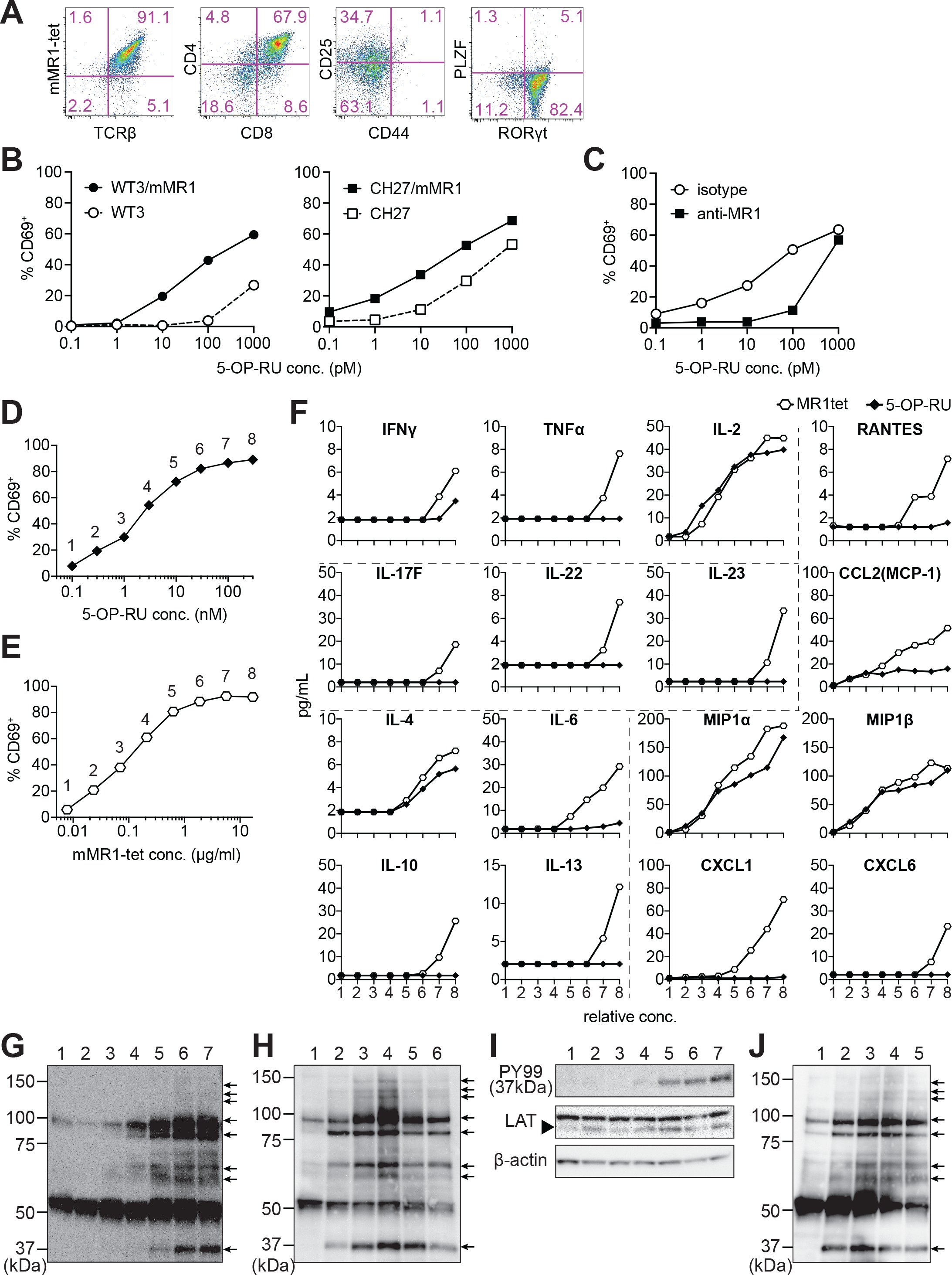
Characterization of m-reMAIT cells. **A. Flow cytometric profiles of m-reMAIT cells** mMR1-tet staining and expression of TCRβ, CD4, CD8, CD25, and CD44 and the transcription factors PLZF and RORγt in m-reMAIT cells on differentiation day 18. **B. 5-OP-RU dose-dependent activation of m-reMAIT cells** The percentages of CD69^+^ cells among m-reMAIT cells challenged with the indicated concentration of 5-OP-RU in the presence of WT3 (○), WT3/mMR1(●), CH27(□), and CH27/mMR1(▪). **C. MR1-dependent activation of m-reMAIT cells** The percentage of CD69^+^ cells among m-reMAIT cells cultured with CH27/mMR1, challenged as in B in the presence of the anti-MR1 antibody (▪) or the isotype control antibody (○) **D. 5-OP-RU dose-dependent activation** The percentage of m-reMAIT cells expressing CD69 upon a challenge with various concentrations of 5-OP-RU. Representative data from two independent experiments are shown. **E. mMR1-tet dose-dependent activation** The percentage of m-reMAIT cells expressing CD69 upon a challenge with the indicated amounts of mMR1-tet. Representative data from two independent experiments are shown. **F. 5-OP-RU- and mMR1-tet-induced cytokines and chemokines** m-reMAIT cells were stimulated with various concentrations of 5-OP-RU (○) or mMR1-tet (▪) and the resultant cytokines and chemokines were quantified with LegendPlex. The concentrations at which each reagent induced a similar degree of activation (% CD69) are shown as relative concentrations (0.1 to 100 nM for 5-OP-RU and 0.01 to 10 μg/ml for mMR1-tet). The number on the X-axis corresponds to that in D and E. **G. Tyrosine phosphorylation elicited with 5-OP-RU** A Western blot analysis with PY99 (anti-phosphotyrosine). Upon a challenge with different concentrations of 5-OP-RU for 30 min, the cell lysate from m-reMAIT cells was separated on SDS-PAGE (5 × 10^5^/lane), and subjected to Western blotting. Lane 1, 0, lane 2, 0.1, lane 3, 1.0, lane 4, 10, lane 5, 100, lane 6 1,000, and lane 7 10,000 (nM) Phosphorylated proteins are indicated with arrows. **H. Time course of tyrosine phosphorylation** A Western blot analysis with PY99. The cell lysate from m-reMAIT cells challenged with 100 nM of 5-OP-RU for the indicated time was separated on SDS-PAGE (5 × 10^5^/lane) and subjected to Western blotting. Lane 1, 0, lane 2, 15, lane 3, 30, lane 4, 60, lane 5, 150, lane 6, 300 (min) Arrows indicate phosphorylated proteins. **I. LAT as a phosphorylated 37-kD protein** A Western blot analysis with PY99, anti-LAT, and anti-β-actin. The cell lysate prepared as described in H for 60 min was subjected to Western blotting. A blot with PY99 (upper panel), anti-LAT (middle panel), and anti-β-actin (lower panel). Phosphorylated LAT and LAT as well as β-actin are indicated (arrow). Lane 1, 0, lane 2, 0.1, lane 3, 1.0, lane 4, 10, lane 5, 100, lane 6 1000, lane 7 10,000 (nM) **J. Tyrosine phosphorylation induced by mMR1-tet** A Western blot analysis with PY99. The cell lysate from m-reMAIT cells challenged with the indicated amounts of unlabeled mMR1-tet for 60 min was subjected to Western blotting (5 × 10^5^/lane). Lane 1, 0, lane 2, 0.43, lane 3, 1.3, lane 4, 4.3, lane 5, 13 (μg/ml) Arrows indicate phosphorylated proteins.

m-reMAIT cells were activated in a manner that was dependent on both 5-OP-RU and MR1 in the presence of APCs, such as WT3, the mouse embryonic fibroblast cell line, WT3/mMR1, WT3-overexpressing murine MR1, CH27, a B1B lymphoma cell line expressing a low level of endogenous MR1 on the cell surface, and CH27/mMR1, CH27-overexpressing murine MR1 (Cheng et al, 1999, Huang et al, 2008, Miley et al, 2003). Similarly, 5-OP-RU dose-dependent activation was observed in the other m-reMAIT cell lines L3-1, L11-1, L15-1, and L19-1 (Figure S1I). The activation of m-reMAIT cells in the presence of CH27 or CH27/mMR1 in turn induced the production of a battery of inflammatory cytokines and chemokines, such as IFN-γ, TNF-α, IL-17A, MIP1β (CCL4), and RANTES (CCL5), as well as IL-2 and the T helper (Th)2 cytokine IL-13, in a 5-OP-RU dose-dependent manner (Figure S1J). 5-OP-RU and mMR1-tet each activated m-reMAIT cells, and promoted the production of a similar, but not identical, set of cytokines and chemokines in a dose-dependent manner in the absence of APCs. It is important to note that mMR1-tet was superior to 5-OP-RU at inducing TNF-α, Th2 cytokines, such as IL-6, IL-10, and IL-13, the Th17 cytokines, IL-17F, IL-22, and IL-23, and inflammatory chemokines, such as RANTES (CCL5), MCP1 (CCL2), CXCL1, and CXCL6, irrespective of their similar abilities to up-regulate CD69 (Figures 1D-F).

Although a TCR stimulation elicits an array of signaling cascades, including protein-tyrosine kinases, phosphatases, GTP-binding proteins, and adaptor proteins, we herein focused on whether 5-OP-RU activated protein-tyrosine kinases and induced tyrosine phosphorylation. The results obtained showed the dose- and time-dependent phosphorylation of proteins (Figures 1G-H). Linker for the activation of T cells (LAT) was identified as a TCR signaling component among phosphorylated proteins (Figure 1I). Unexpectedly, the addition of mMR1-tet, a reagent widely used to detect MAIT cells, engendered similar protein tyrosine phosphorylation, as was the case for 5-OP-RU (Figures 1G-H and 1J).

These results demonstrated that MAIT-iPSCs gave rise to m-reMAIT cells upon differentiation under T-cell permissive conditions, and m-reMAIT cells were activated by 5-OP-RU and mMR1-tet in the absence of APCs concomitant with protein tyrosine phosphorylation and the production of cytokines and chemokines.

### m-reMAIT cell migration into different organs

To follow the dynamics of m-reMAIT cells, cells were adoptively transferred into mice and their migration and maturation status were examined. To distinguish donor cells from endogenous MAIT cells, C57BL/6 (Ly5.1) mice received m-reMAIT cells that were Ly5.2. m-reMAIT cells and endogenous MAIT cells were detected in the thymus, bone marrow, lung, liver, spleen, intestines, and the mediastinal and inguinal lymph nodes at various endogenous MAIT cells/m-reMAIT cell ratios (Figure 2A).

**Figure 2.**
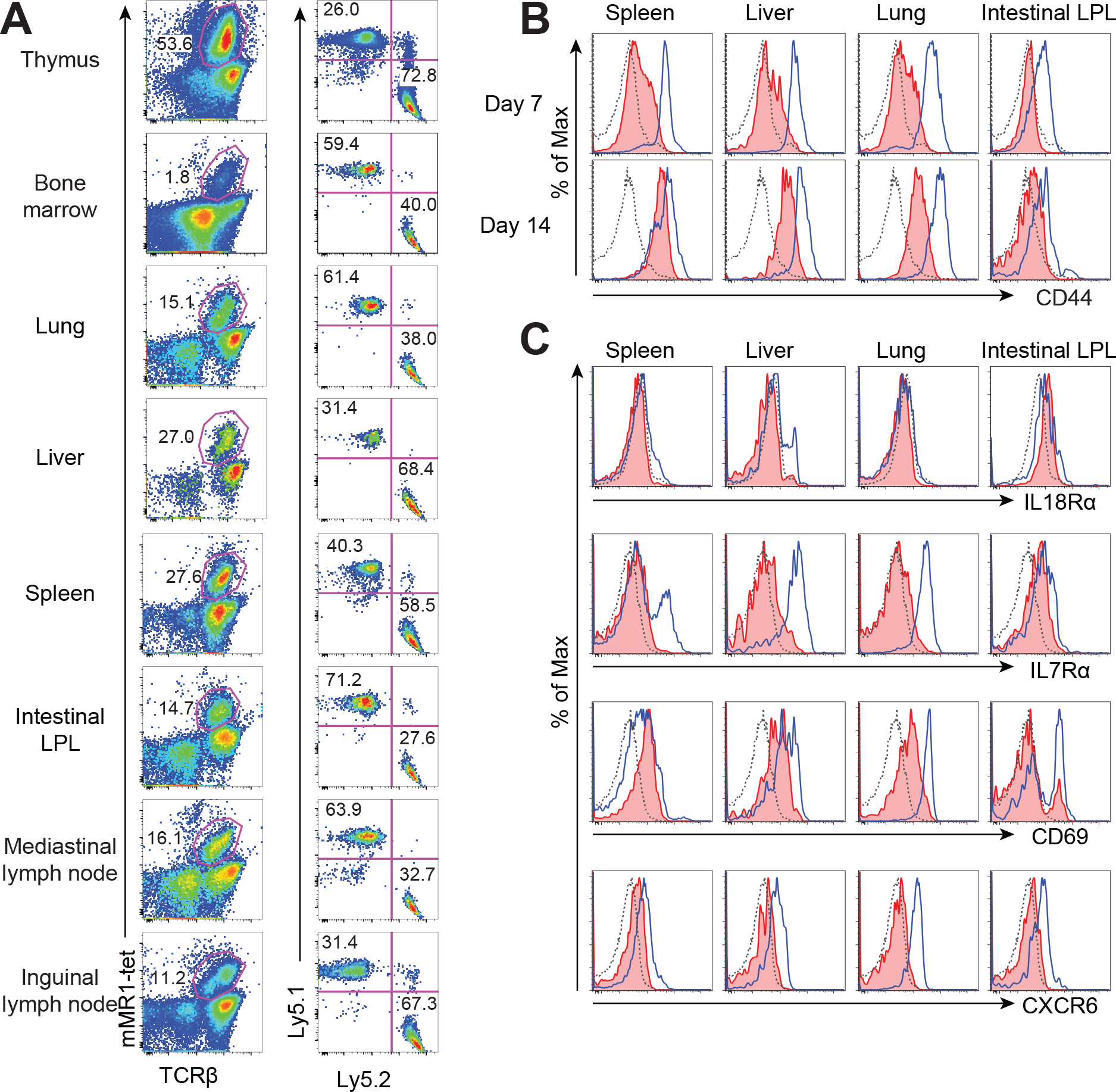
Behavior of m-reMAIT cells upon adoptive transfer. **A. m-reMAIT cell migration into different organs** m-reMAIT cells (Ly5.2) were adoptively transferred into C57BL/6 (Ly5.1) mice via an intraperitoneal injection (1 × 10^7^ cells/mouse), and endogenous as well as exogenous TCRβ^+^mMR1-tet^+^ cells were enriched 7 days later with mMR1-tet (left panel) and subjected to analyses of the expression of Ly5.1 and Ly5.2 (right panel). The number in the panel shows the percentage of Ly5.1 (endogenous) and Ly5.2 (exogenous) TCRβ^+^mMR1-tet^+^ cells. Representative data from a pool of 3-4 mice per experiment are shown. **B. Time-dependent up-regulation of CD44** CD44 expression in TCRβ^+^mMR1-tet^+^ cells from the indicated tissues 7 and 14 days after m-reMAIT cell adoptive transfer. Isotype control (dotted line), endogenous MAIT cells (Ly5.1^+^TCRβ^+^mMR1-tet^+^ cells) (plain blue line), and exogenous m-reMAIT cells ((Ly5.2^+^TCRβ^+^mMR1-tet^+^ cells) (shaded in red). Representative data from a pool of 3-4 mice per experiment are shown. **C. Expression of molecules relevant to MAIT cells** The expression of molecules in m-reMAIT cells from the indicated tissues 14 days after adoptive transfer. Isotype control (dotted line), endogenous MAIT cells (Ly5.1^+^TCRβ^+^mMR1-tet^+^ cells) (plain blue line), and exogenous m-reMAIT cells (Ly5.2^+^TCRβ^+^mMR1-tet^+^ cells) (shaded in red). Representative data from a pool of 3-4 mice per experiment are shown.

Since CD44 levels reflect the degree of the functional maturation of T cells, we examined time-dependent changes in CD44 expression in m-reMAIT cells (Budd et al, 1987). In comparisons with endogenous MAIT cells, m-reMAIT cells showed the time-dependent up-regulation of CD44 in the spleen, liver, and lungs, and weaker expression in the intestines (Figure 2B).

An analysis of other molecules relevant to the functionality of MAIT cells revealed quasi-equivalent IL-18Rα expression in the spleen, liver, and intestines to that in endogenous cells. Moreover, the expression of IL-7Rα and CXCR6 in all organs examined was weaker than in endogenous cells, while CD69 expression was quasi-equivalent to that in endogenous cells (Figure 2C).

These results indicated that m-reMAIT cells migrated into different organs accompanying maturation upon adoptive transfer in immunocompetent mice.

### Tumor inhibitory activity of m-reMAIT cells

Since the role of MAIT cells in tumors has been a target for intensive scrutiny, we attempted to clarify whether m-reMAIT cells interfere with tumor metastasis and improve mouse survival (Godfrey et al, 2019, Toubal et al, 2019). A previous study on B16F10 melanoma suggested that MAIT cells induce tumor development in a manner that is dependent on MR1 and agonists, such as 5-OP-RU (Yan et al, 2020). While B16F10 significantly up-regulated MR1 on the cell surface with the 5-OP-RU challenge, similar changes were not observed for Lewis lung carcinoma (LLC) (Figure 3A). Therefore, the use of LLC allows for assessments of the role of m-reMAIT cells in tumor immunosurveillance independent of the 5-OP-RU-MR1 axis.

**Figure 3.**
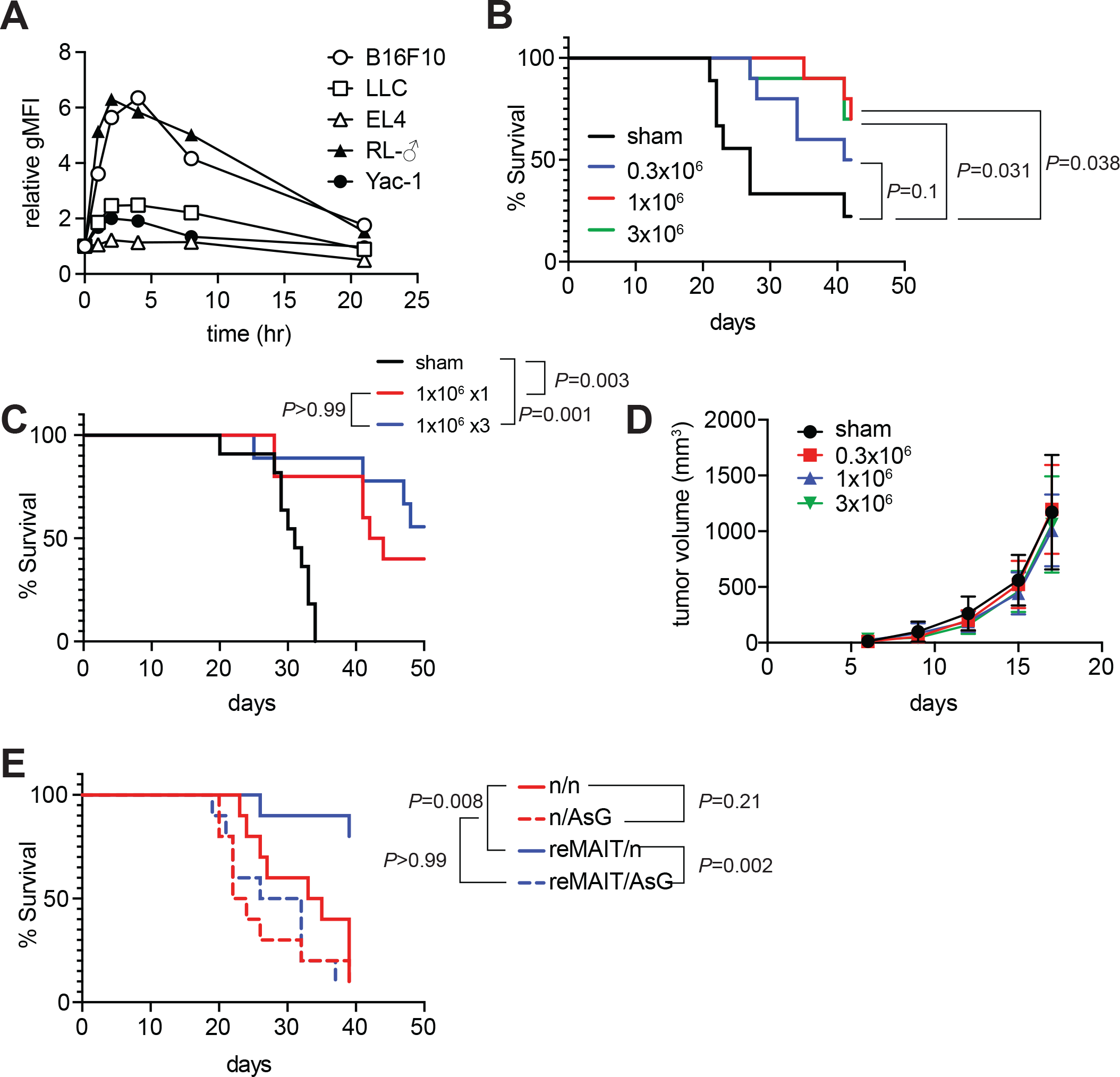
Extensions in mouse survival by m-reMAIT cells. **A. Time course of 5-OP-RU-dependent MR1 expression** The indicated cancer cell lines were challenged with 5-OP-RU. MR1 expression levels on the cell surface at the indicated time point are shown as relative gMFI. Data are representative of three independent experiments. **B.** m-reMAIT cell dose-dependent survival extension C57BL/6 mice received the indicated amounts of m-reMAIT cells 6 days prior to the LLC inoculation (3 × 10^5^ cells/mouse i.v.), and survival was monitored (n=10∼12/group). Data are representative of three independent experiments. *P* values between the indicated group are shown (the Log-rank test). **C.** Effects of the multiple transfer of m-reMAIT cells on survival The survival of C57BL/6 mice that received m-reMAIT cells (1 × 10^6^/mouse, i.p.) 6 days prior to the LLC inoculation (3 × 10^5^ cells/mouse, i.v.), and of mice that received LLC and two more consecutive transfers of m-reMAIT cells (1 × 10^6^/transfer/mouse) was monitored (n=10∼12/group). Sham-treated mice that only received LLC served as a control. Data are representative of two independent experiments. *P* values between the indicated groups are shown (the Log-rank test). **D.** Effects of m-reMAIT cells on *in situ* tumor growth Growth curve of LLC. LLC (3 × 10^5^/mouse) were subcutaneously inoculated into the right flank of C57BL/6 mice 6 days after the m-reMAIT cell transfer (i.p.). Tumor size was plotted with time. Sham treated (●), 0.3 × 10^6^ transferred (▪), 1.0 × 10^6^ transferred (▴), and 3.0 × 10^6^ m-reMAIT cells transferred (▾). Data are shown with SEM (5-6 mice per group).

The adoptive transfer of m-reMAIT cells into mice followed by the LLC inoculation resulted in the extension of survival in a dose-dependent manner. While 3 × 10^5^ m-reMAIT cells slightly extended survival over the control, more than 1 × 10^6^ m-reMAIT cells significantly extended survival (Figure 3B). Nevertheless, multiple transfers of m-reMAIT cells failed to confer significant extensions in survival (Figure 3C). In contrast to the above findings, the adoptive transfer of m-reMAIT cells failed to suppress tumor growth at any dose following the subcutaneous inoculation of LLCs, which represents an *in situ* tumor growth model (Figure 3D).

These results implied that m-reMAIT cells inhibited tumor metastasis rather than suppressing tumor growth *in situ*.

### Cytolytic activity of m-reMAIT cells in combination with NK cells

While the above results indicated that m-reMAIT cells functioned to suppress metastasis, it currently remains unclear whether other immune cells are involved in this tumor inhibitory activity. Since NK cells are an essential innate sentinel in tumor immunosurveillance, we speculated that NK cells and m-reMAIT cells may cooperate in tumor immunity. To investigate the interaction between NK cells and m-reMAIT cells, they were cocultured and their impact was assessed. While m-reMAIT cells were activated in a NK cell dose-dependent manner, the activation of NK cells was less marked (Figure 4A). Moreover, a coculture led to the production of Th1 and Th17 cytokines as well as inflammatory chemokines in a NK cell dose-dependent manner (Figure 4B).

**Figure 4.**
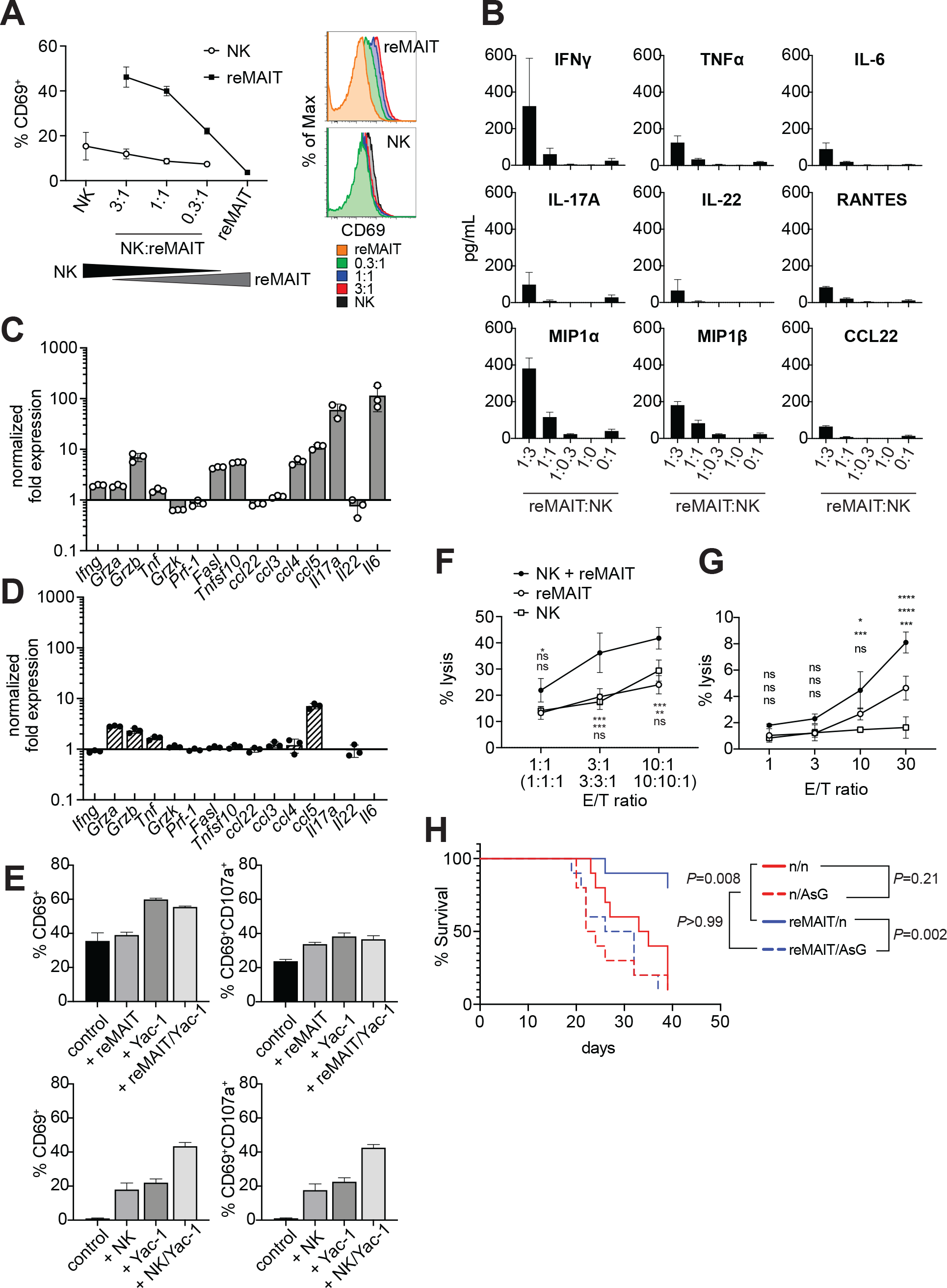
Anti-tumor activity of m-reMAIT cells bolstered by NK cells. **A. Activation of m-reMAIT cells by NK cells** CD69 expression in m-reMAIT cells (reMAIT ●) and NK cells (NK ◯) upon an incubation at the indicated ratio. The percentage of cells expressing CD69 (left panel) and the intensity of CD69 in each cell population (right panel) are shown. **B. Cytokines and chemokines upon a coculture** Cytokines and chemokines released upon a coculture of m-reMAIT cells and NK cells at the indicated ratio are shown (reMAIT:NK). Amounts were quantified with LegendPlex. Representative data from two independent experiments are shown. **C. Transcripts relevant to cytolytic activity in m-reMAIT cells** *Ifng*, *Gzma*, *Gzmb*, *Tbf*, *Gzmk*, *Pfr1*, *Fasl*, *Tnfsf10*, *Il6*, *Il17a*, *Il22*, *Ccl3*, *Ccl4*, *Ccl5*, and *Ccl22* in m-reMAIT cells cultured with NK cells were quantified with qRT-PCR. m-reMAIT cells and NK cells were sort-purified after the coculture (purity >98%) or cultured individually. The expression of each transcript was normalized with *Gapdh* and fold changes in the relative expression of the transcript in m-reMAIT cells cultured with NK cells relative to that in m-reMAIT cells cultured alone are shown. Data are representative of three independent experiments. **D. Transcripts relevant to cytolytic activity in NK cells** Fold changes in the relative expression of the indicated transcript as described in C in NK cells cultured with m-reMAIT cells relative to that in NK cells alone are shown. Representative data from three independent experiments are shown. **E. Activation and degranulation of NK cells and m-reMAIT cells** The expression of CD69, an activation marker, and CD107a, a marker for the exocytosis of cytolytic granules, was assessed under various culture conditions. The percentages of CD69^+^ cells and CD69^+^CD107a^+^ cells among NK cells alone (control), NK cells cocultured with m-reMAIT cells (+reMAIT), NK cells cultured with Yac-1 (+Yac-1), and NK cells cocultured with m-reMAIT cells and Yac-1 (+reMAIT/Yac-1) (upper panels). The percentages of CD69^+^ cells and CD69^+^CD107a^+^ cells among m-reMAIT cells alone (control), m-reMAIT cells cocultured with NK cells (+NK), m-reMAIT cells cocultured with Yac-1 (+Yac-1), and m-reMAIT cells cocultured with NK cells and Yac-1 (+NK/Yac-1) (lower panels). Data are representative of three independent experiments. **F. Cytolytic activity against Yac-1** Cytolytic activity of m-reMAIT cells (reMAIT ◯), NK cells (NK ☐), and NK cells plus m-reMAIT cells (NK+reMAIT ●). Cytolytic activities (% lysis) at different effector (NK cells, m-reMAIT cells, and NK cell + m-reMAIT cells)/Target (Yac-1) (E/T) ratios are shown. Representative data from three experiments are shown. The significance of differences between the groups at the indicated E/T ratio assessed with a two-way ANOVA is shown (* P<0.05, \*\**P*<0.01, \*\*\**P*<0.005). From the top, NK+reMAIT vs. reMAIT, NK+reMAIT vs. NK, and reMAIT vs. NK. Data are representative of three independent experiments. **G. Cytolytic activity against LLC** The cytolytic activities of m-reMAIT cells (◯), NK cells (☐), and m-reMAIT cells plus NK cells (●) against LLC at the indicated E/T ratio are shown as % lysis. The significance of differences between the groups is calculated as in E. Data are representative of three independent experiments. **H. NK cell-dependent extension of survival** C57BL/6 mice were divided into two groups, one that received 1 × 10^6^ m-reMAIT cells (reMAIT) and another that was left untreated (n). Each group was further divided into two subgroups, one that received consecutive injections of anti-Asialo GM1 (AsG) (-1 and 16 days, 50 μg/mouse) after the LLC inoculation (3×10^5^ I.V.) and another that was left untreated (n). Survival was monitored thereafter. Representative data from two independent experiments are shown (n=10∼14/group). *P* values between the indicated groups are shown (the Log-rank test).

To obtain further insights into the molecular events underlying this interaction, NK cells and m-reMAIT cells were purified with FACS after the coculture and each subset was examined for transcripts relevant to cytolytic function. *Ifng* was induced in m-reMAIT cells, but not in NK cells. Similarly, the coculture enhanced the transcripts for serine proteases, such as *Grza*, *Grzb*, and *Tnf*, in m-reMAIT cells and NK cells (Trapani & Smyth, 2002). In contrast, the expression of the transcripts for the TNF superfamily, including *Fasl* and *Tnsf10*, was only stimulated in m-reMAIT cells. Moreover, the expression of *Mip1b (Ccl4)*, *Rantes (Ccl5)*, *Il17a*, and *Il6* increased in reMAIT cells, whereas only *Rantes (Ccl5)* was up-regulated in NK cells (Figures 4C-D).

We then assessed whether this change had an impact on CD69 and CD107a, a marker of cytolytic granule exocytosis that is a prerequisite for cytolytic activity. While the coculture with m-reMAIT cells did not markedly up-regulate the expression of CD69 (CD69^+^ cells) in NK cells, the addition of Yac-1 (alone or in combination with m-reMAIT cells) increased this percentage. However, these combinations resulted in negligible changes in the percentage of CD69^+^CD107a^+^ cells (Figure 4 E, upper panels). In contrast, an incubation with NK cells or Yac-1 enhanced the percentage of CD69^+^ cells in m-reMAIT cells. Moreover, a coculture with NK cells together with Yac-1 resulted in further increases in this percentage, and similar results were obtained for CD69^+^CD107a^+^ cells in m-reMAIT cells (Figure 4 E, lower panels).

We then investigated whether the above activation and/or expression of CD107a reflected lytic activity against tumor cells. m-reMAIT cells lysed Yac-1, a NK cell-sensitive tumor cell line, as efficaciously as NK cells, and these cells together enhanced killing (Figure 4F). We extended our study to LLC, a NK cell-insensitive cell line. NK cells alone did not induce cell death in LLC, whereas m-reMAIT cells did. Moreover, the combination of these cells exhibited synergistically enhanced lytic activity (Figure 4G).

We also investigated the significance of the interaction between NK cells and m-reMAITs cell *in vivo*. To highlight the role of NK cells in m-reMAIT cell-mediated anti-metastasis activity, NK cells were depleted from mice and the impact of this change was assessed. The depletion of NK cells with an anti-asialo GM1 antibody (AsG) did not significantly affect the survival of control mice. However, the AsG treatment abrogated the survival advantage of m-reMAIT cells, highlighting the stimulatory role of NK cells in m-reMAIT cell-mediated tumor inhibitory activity (Figure 4H).

These results demonstrated that the intrinsic cytolytic activity of m-reMAIT cells against tumors was bolstered by NK cells concomitant with the up-regulation of the relevant arsenals.

## Discussion

In the present study, we demonstrated that the presence of rearranged TCR loci specific to MAIT cells in the alleles constrained the fate of pluripotent cells when they differentiated into the T-cell lineage, which is consistent with our previous findings on human MAIT-like cells (Wakao et al, 2013). A similar phenomenon was observed for iNKT-like cell differentiation from cloned embryonic stem (ES) cells (Wakao et al, 2008). Since iNKT cells and MAIT cells belong to a family of innate-like T cells harboring semi-invariant TCR, the expression of the machinery relevant to TCR rearrangements may be repressed during differentiation *in vitro*. In accordance with this notion, transcripts for the machinery, such as *Rag1*, *Rag2*, and *Dntt*, were found to be repressed in human MAIT-iPSCs concomitant with epigenetic silencing upon differentiation (Saito et al, 2017). Therefore, further studies are warranted to clarify whether the same corollary applies to the murine system.

It is important to note that m-reMAIT cells were competent in TCR signaling. The present result showing the activation of m-reMAIT cells and subsequent production of cytokines and chemokines in the absence of APCs suggested that 5-OP-RU directly operated via MR1 in m-reMAIT cells (Figures 1D-F). 5-OP-RU may be passively transported into cells, which in turn promotes the release of antigen-loaded MR1 from the endoplasmic reticulum (ER) onto the plasma membrane, thereby enabling signaling via TCR in other m-reMAIT cells (referred to as a cis-acting signal) regardless of the apparent absence of MR1 on the cell surface (Figures 1B, 1D, and S1I) (McWilliam et al, 2016). Therefore, in the absence of APC, a markedly higher concentration of 5-OP-RU may be required to activate m-reMAIT cells than that in the presence of APC.

In contrast to 5-OP-RU, mMR1-tet may directly bind to TCR on m-reMAIT cells and elicit intracellular signals (referred to as a trans-acting signal). Therefore, it is important to elucidate differences in the 5-OP-RU-emenated cis-acting signal and trans-acting signal elicited by mMR1-tet. Although the cis- and trans-acting signals both induced a similar set of tyrosine-phosphorylated proteins, they differed in their ability to produce cytokines and chemokines (Figures 1F-H and 1J). Therefore, 5-OP-RU and mMR1-tet may have activated different signals, leading to differences in the pattern of serine/threonine phosphorylation or dephosphorylation via TCR, and ultima tely a different profile of cytokine and chemokine production (Gaud et al, 2018, Nausch & Cerwenka, 2008). In this respect, the signaling pathway(s) responsible for the production of IL-2, IL-4, MIP1α (CCL3), and MIP1β (CCL4) may be common between 5-OP-RU and mMR1-tet.

While 5-OP-RU induced protein tyrosine phosphorylation, the identity of these proteins remains unclear, except for LAT (Figures 1G-H). Since LAT forms a hub that interacts with many signaling molecules, such as transmembrane receptors, phosphatases, kinases, guanine nucleotide exchange factors, GTP-activating proteins, ion channels, transporters, and ubiquitin ligases, further studies will provide insights into the proteins responsible for MAIT-TCR signaling and reveal differences in signaling between conventional T cells and MAIT cells (Malissen et al, 2014).

It is important to note that the challenge with mMR1-tet induced the activation of m-reMAIT cells and production of cytokines and chemokines. This result suggests that MAIT cells prepared from animals, including humans and mice, are unintentionally stimulated during experiments. Therefore, the interpretation of data requires caution (Figures 1E-F and 1J).

Although a previous study suggested that MAIT cells comprise MAIT1 and MAIT17 subsets characterized by the Th-1 and Th-17 transcriptome, respectively, the present results revealed that m-reMAIT cells produced Th-1, Th-2, and Th-17 cytokines (Salou et al, 2019). This may reflect a developmental stage at which m-reMAIT cells are en route to functional maturation and m-reMAIT cells generated *in vitro* represent the most immature stage. Therefore, it is tempting to postulate that the interaction between nascent MAIT cells and double-positive thymocytes favors the differentiation of naïve MAIT cells into MAIT1 and MAIT17 *in vivo*.

The results on the adoptive transfer of m-reMAIT cells indicated that nascent MAIT cells egressed from the thymus migrate into any organs concomitant with the up-regulation of CD44, a cell adhesion receptor, memory, and/or activation marker (Figure 2B) (Benlagha et al, 2005). It is important to note that the expression of IL-7Rα in m-reMAIT cells became quasi-equivalent to that in endogenous cells in the intestines (Figure 2C). Since IL-7 from intestinal epithelial cells (IEC) play a pivotal role in the homeostasis of IEC, the up-regulation of IL-7Rα in m-reMAIT cells may mirror an intrinsic and primary role in this homeostasis and/or in the integrity of the intestinal epithelial barrier (Rouxel et al, 2017, Shalapour et al, 2012). Alternatively, the up-regulation of IL-7Rα may reflect the role of MAIT cells in IL-17A and Th1 cytokine production, which is dependent on IL-7 (Tang et al, 2013). Together with the time-dependent up-regulation of CD44, increases in IL-18Rα and CXCR6 in different organs indicated that m-reMAIT cells mature in the host, which has important implications for cell therapy.

m-reMAIT cell dose-dependent mouse survival indicated their tumor inhibitory activity and function as effector cells in tumor surveillance (Figure 3B).

While the importance of NK cells in tumor surveillance is well known, that of MAIT cells remains elusive (Guillerey et al, 2016, Sharma et al, 2017). The present results showing the NK cell-dependent activation of m-reMAIT cells and the subsequent production of inflammatory cytokines and chemokines suggested that NK cells stimulated the tumor inhibitory effects of MAIT cells (Figures 4A-B). Moreover, the enhanced expression of transcripts relevant to the cytolytic function of m-reMAIT cells upon an interaction with NK cells *in vitro* indicated that m-reMAIT cells exhibit cytolytic activity *in vivo* (Figures 4C-G). The NK cell-stimulated up-regulation of *Grzb*, *Fasl*, and *Tnfsf10* in m-reMAIT cells suggested that cytolytic activity comprised exocytosis-mediated killing with GrzB, a serine protease, and FasL- and Tnfsf10-mediated killing through the activation of caspases (Rossin et al, 2019, Trapani & Smyth, 2002). The present result showing that NK cell depletion compromised the survival of mice that had received m-reMAIT cells upon a tumor inoculation is consistent with the above findings, further highlighting the role of NK cells in boosting the tumor inhibitory activity of m-reMAIT cells *in vivo*. While NK cells recognize NKG2D ligands and/or the lack of MHC I on tumor cells and eradicate them, the molecular mechanisms by which m-reMAIT cells recognize and eliminate tumor cells with the aid of NK cells warrants further study (Nausch & Cerwenka, 2008).

In contrast to the present results, MAIT cells have been shown to induce the metastasis of melanoma B16F10 (Yan et al, 2020). Although the reason for this discrepancy currently remains unclear, differences in the dynamics of MR1 shuttling between ER and the plasma membrane upon a 5-OP-RU challenge or putative ligand(s) present in the tumor milieu (TME) may be a key feature. Since LLC did not strongly up-regulate MR1 on the cell surface, in contrast to B16F10, when stimulated (Figure 4A), 5-OP-RU may have bolstered B16F10 tumorigenicity through MR1. The MR1-elicited signal may suppress the function of NK cells for B16F10, whereas signal(s) from putative ligand(s) present in TME via MR1 did not, thereby enhancing or preserving NK cell function. Further studies are needed to elucidate the underlying mechanisms, which will provide a novel avenue for tumor immunotherapy with iPSC-derived MAIT cells.

Although we demonstrated that m-reMAIT cells exhibited cytolytic activity against LLC together with NK cells *in vitro* (Figures 4C and 4E-G), difficulties are associated with demonstrating that m-reMAIT cells isolated from mice that had received the cells exhibit similar lytic activity *ex vivo*. This is due to the low recovery of m-reMAIT cells and to the compulsory use of mMR1-tet in cell preparations, which may interfere with cytolytic activity. Furthermore, it currently remains unclear whether these results are applicable to other cancer cells, such as B16F10, if the use of 5-OP-RU enhances or inhibits the cytolytic activity of m-reMAIT cells, and also whether MAIT-like cells prepared from human MAIT-iPSCs exhibit similar activity.

In summary, the adoptive transfer model with m-reMAIT cells used in the present study opens a new avenue for exploiting the function of MAIT cells and providing insights into their interaction(s) with immune cells in immunity.

## Materials and Methods

### Mice

All mouse experiments were performed with approval from the Institutional Animal Care and Use Committee of Dokkyo Medical University. C57BL/6N mice were purchased from CLEA Japan (Tokyo, Japan). C57BL/6 (Ly5.1) mice were obtained from the RIKEN Bioresource Center and bred in-house. All mice were housed in the Animal Research Center, Dokkyo Medical University, under specific pathogen-free conditions with controlled lighting and temperature with food and water provided *ad libitum*. Male C57BL/6N mice aged 6 weeks were used to isolate MAIT cells for the generation of iPSCs. Male and female mice aged between 8 to 12 weeks were used in adoptive transfer experiments and in tumor experiments.

### Cell lines

OP9/DLL1 cells were maintained in αMEM supplemented with 20% FBS. The mouse cancer cell lines B16F10, CH27, CH27/mMR1, EL4, and LLC were cultured in DMEM supplemented with 10% FBS, while RL-♂1, WT3, WT3/mMR1 and Yac-1 were cultured in RPMI1640 supplemented with 10% FBS at 37°C in 5% CO_2_.

### Antibodies

The antibodies used in the present study are listed in the supplemetal table 2.

### Oligonucleotides

The oligonucleotides used in the present study are summarized in the oligonucleotide Table.

### Preparation of mouse immune cells

#### Spleen, thymus, and lymph nodes

Tissues were prepared by mashing through a 40-µm mesh cell strainer with a syringe plunger. Single cells were suspended in RPMI1640 supplemented with 10% FBS, 10 mM HEPES pH 7.0, 0.1 mM 2-mercaptoethanol, and 100 IU/ml of penicillin/streptomycin (referred cR10) and spun down at 400×*g* for 4 min. To lyse red blood cells, the cell pellet was suspended in autoclaved ice-cold MilliQ water for 15 sec and immediately neutralized with 4% FBS in 2× PBS. After centrifugation, cells were resuspended in cR10.

#### Lungs and liver

Single-cell suspensions from the lungs and liver were prepared using enzymatic digestion. Briefly, tissues were placed into a GentleMACS C-tube (Miltenyi Biotec) and cut into approximately 5-mm^3^ pieces. Four milliliters of tissue digestion solution (90 U/ml collagenase Yakult, 275 U/ml collagenase type II, 145 PU/ml Dispase II, and 4% BSA in HBSS) was added per tissue and tissues were homogenized using the GentleMACS dissociator (Miltenyi Biotech) with the following program: m_lung_01_02 for the lungs and m_liver_03_01 for the liver. Suspensions were then incubated at 37°C for 30 min under gentle rotation, followed by dissociation with m_lung_02_01 for the lungs and m_liver_04_01 for the liver, and subjected to discontinuous density centrifugation over layers of 40% and 60% Percoll at 400×*g* for 20 min. Cells were recovered from the 40%-60% Percoll interface, washed with PBS, and then suspended in cR10.

#### Intestines

The intestines were longitudinally incised and their contents were thoroughly washed out three times by vigorous shaking. Tissue dissected into 1-cm pieces were placed into a 50-ml conical tube and washed vigorously by shaking three times with PBS. After discarding the supernatant, tissues were treated with 40 ml of intraepithelial lymphocyte (IEL)-washing solution (HBSS containing 1 mM DTT, 5 mM EDTA, and 1% BSA) by shaking vigorously at 37°C for 30 min under gentle rotation. After washing three times with MACS buffer (PBS containing 2 mM EDTA and 0.5 % BSA), tissue was washed again with 40 ml HBSS. After removal of the supernatant, tissues were placed into a GentleMACS C-tube (Miltenyi Biotec), cut into small pieces with scissors, and 4 ml of the Tissue digestion solution (see above) was added. Tissues were processed using the following program: m_brain_01_02 and digested at 37°C for 30 min under gentle rotation followed by dissociation with the program: m_intestine_01_01. Cell suspensions were subjected to Percoll discontinuous density centrifugation and isolated cells were resuspended in cR10.

### Generation of iPSCs from mouse MAIT cells

Single-cell suspensions from the lungs were stained with APC-labeled 5-OP-RU-loaded mouse MR1 tetramer (mMR1-tet) (NIH Tetramer Core Facility) before magnetic enrichment with anti-APC microbeads and LS or MS columns (Miltenyi Biotech) according to the manufacturer’s instructions. Enriched cells were stained with FITC-CD44, PE-B220, PE-F4/80, and PE/Cy7-TCRβ and MAIT cells were then sorted as B220^−^F4/80^−^TCRβ^+^CD44^hi^mMR1-tet^+^ cells with the FACSJazz cell sorter (BD Biosciences). Regarding transduction, purified MAIT cells (1800 – 6300 cells) were placed into the wells of 96-well plastic plates coated with anti-CD3 and anti-CD28 antibodies (15 and 20 µg/ml, respectively) for 18 h and then infected with Sendai virus KOSM302L at MOI of 25 at 37°C in 5% CO_2_ under gentle orbital shaking for 2.75 hrs. After transduction, virus-containing medium was removed by centrifugation at 400×*g* for 4 min. Cells were refed with mouse ES medium containing StemSure D-MEM (FUJIFILM Wako), 15% FBS (BioSera), 1× nonessential amino acids (FUJIFILM Wako), 2 mM L-glutamine (Nacalai Tesque), 0.1 mM 2-mercaptoethanol (FUJIFILM Wako), 100 U penicillin/100 µg streptomycin (Lonza), 1000 U/ml mouse LIF (FUJIFILM Wako), and transferred to the wells of 6-well culture plates seeded with mitomycin C (MMC)-treated MEF. Medium was changed every 2-3 days. Three to four weeks after infection, ES-like colonies were picked up and treated with 0.25% Trypsin-1 mM EDTA (TE) at 37°C for 10 min, and individually transferred to the wells of 24-well culture dishes seeded with MMC-treated MEF. Among 46 iPSCs isolated after reprogramming, five iPSCs (L3, L7, L11, L15, and L19) were subjected to limiting dilutions after 2-3 passages and a monoclonal iPSC line was generated and designated accordingly (e.g., L3-1, L7-1∼8, L11-1∼4, L15-1∼3, and L19-1∼6) (Figures S1B and S1D). The established iPSC lines were maintained with mouse ES medium supplemented with 3 µM CHIR99021 (FUJIFILM Wako) and 10 µM PD0325901 (FUJIFILM Wako). The majority of experiments were performed with L7-1, except for the PCR analysis in Figure S1A, in which samples were used before limiting dilutions. Monoclonal iPSC lines were cryopreserved with serum-free cell culture freezing medium BAMBANKER (GC lymphotec) in liquid nitrogen.

### PCR detection of the rearranged configuration of *TCR* loci in MAIT-iPSCs

To confirm that MAIT-iPSCs stemmed from MAIT cells, PCR detecting the rearranged configuration of *TRAV* specific for MAIT cells was performed with the primer sets ADV19 and AJ33. Genomic DNA was prepared from MAIT-iPSCs with NaOH. To detect *TRBV* in MAIT-iPSCs, total RNA was prepared from m-reMAIT cells (days 20-28 of differentiation, varying according to the clones), using the RNeasy Mini kit (Qiagen). cDNA was synthesized with the First strand cDNA synthesis kit (Thermo Fisher Scientific) and subjected to PCR with the primer sets TRBV13 and TRBC-Rev, and TRBV9 and TRBC-Rev followed by DNA sequencing (Fasmac). The usage of TRBV-D-J was analyzed by IGBLAST (NCBI, NIH https://www.ncbi.nlm.nih.gov/igblast/).

### Southern blot analysis to detect rearranged *TRAV*

Genomic DNA (3 μg) prepared from MAIT-iPSCs (L7-1∼6, L11-1∼4, L15-1∼3, and L19-1∼6) with NucleoSpin Tissue (NACHREY-NAGEL) was digested with 40 U *BamH*1 (TOYOBO) at 37°C for 18 h, separated by 0.9 % agarose gel electrophoresis, and transferred onto Biodyne membranes (PALL Life Sciences). The membrane was pre-hybridized with PerfectHyb (TOYOBO) at 68°C for 60 min and hybridized with an α-^32^P-dCTP-labeled probe at 68°C for 18 h. The probe was amplified with the primer set TRAV19 (forward) and TRAV19 (reverse) using genomic DNA from C57BL/6 as a template and the resultant 350-bp PCR product was radiolabeled with the Random primer DNA labeling kit ver. 2.0. (Takara)(Figure S1D). The membrane was washed with 2× saline sodium citrate (SSC) and then 0.1 × SSC buffer twice at 68°C, exposed to an imaging plate, and the signal was detected with a phosphoimager (Typhoon 7000, GE Healthcare).

### Differentiation of MAIT-iPSCs into MR1 tetramer^+^ m-reMAIT cells

On day 0, 1.2×10^5^ MAIT-iPSCs (L3-1, L7-1(L7), L11-1, L15-1, and L19-1) were seeded on a 10-cm culture dish containing 2-4 days post-confluent OP9/DLL1 in αMEM supplemented with 10% FBS (Corning). On day 3, culture medium was replaced with fresh medium. On day 5, cells were treated with 2 ml TE and 8 ml of αMEM containing 20% FBS (BioSera) was added and then suspended in wells. The dish was incubated at 37°C in 5% CO_2_ for 45 min. Floating cells were collected and passed through a 40-µm mesh cell strainer. After centrifugation, cells were transferred to a new 10-cm culture dish filled with confluent OP9/DLL1 in αMEM containing 10% FBS (Corning) and 5 ng/ml human recombinant FLT3L. On day 8, loosely attached cells (cells containing lymphocyte progenitors) were harvested with gentle pipetting. After centrifugation, cells were transferred to the wells of a 6-well culture plate containing confluent OP9/DLL1in αMEM supplemented with 20% FBS (BioSera), 5 ng/ml human FLT3L, and 1 ng/ml mouse IL-7. On day 10 and later, culture medium was changed every other day (20% FBS containing αMEM supplemented with 5 ng/ml human FLT3L and 1 ng/ml mouse IL-7). Differentiated cells from iPSCs (m-reMAIT cells) were expanded on a new 10-cm culture dish filled with confluent OP9-DLL1 in αMEM supplemented with 20% FBS (BioSera) and mouse IL-7. m-reMAIT cells harvested on days 21-28 were used in the experiments unless otherwise indicated.

### Preparation of 5-OP-RU

An aliquot of 5-A-RU (3.6 mM in DMSO) was incubated with three volumes of 1 mM methylglyoxal (MilliQ water) at 37°C for 30 min and used for assays. The appropriate synthesis of 5-OP-RU was confirmed with LC-MS/MS (Triple Quad 5500, Sciex).

### m-reMAIT cell activation assay

m-reMAIT cells (L3-1, L7-1(L7), L11-1, L15-1, and L19-1) were incubated with the indicated concentration of 5-OP-RU or mMR1-tet at 37°C in 5% CO_2_ for 18 hours in the presence or absence of equal numbers of CH27, CH27/mMR1, WT3, and WT3/mMR1, and then subjected to a CD69 expression analysis by flow cytometry. The culture supernatant was used for cytokine and chemokine quantification (see below). To examine whether the above activation of m-reMAIT cells was dependent on MR1, an anti-MR1 antibody (26.5) or isotype antibody (10 μg/ml) was added 1 h prior to the addition of 5-OP-RU.

### Doubling time estimation with CFSD-SE (CFSE)

m-reMAIT cells (differentiation on days 18-23) were labeled with CFSE (1.0 *μ*M) and cultured on OP9/DLL1 in *α*MEM containing 20% FBS (BioSera) and 1 ng/ml mouse IL-7. CFSE intensity in the cells was monitored daily by flow cytometry with 488 nm excitation and a bandpass filter of 530/30 nm. The proliferation rate of m-reMAIT cells was calculated from a logarithmical growth curve based on the intensity of CFSE.

### Detection of tyrosine-phosphorylated proteins

m-reMAIT cells (TCRβ^+^mMR1-tet^+^ cells, >95% based on the flow cytometric analysis) were suspended in 20% FBS containing αMEM (1.0 × 10^6^/ml) for the indicated time in the absence or presence of varying amounts of 5-OP-RU or unlabeled mMR1-tet. Cells were harvested by centrifugation at 600×*g* and lysed with lysis buffer (50 mM HEPES-NaOH pHb7.4, 150 mM NaCl, 1% (v/v) Triton-X100, 1 mM sodium vanadate, and 5 mM NaF) supplemented with ×1 protease and phosphatase inhibitor cocktail (FUJIFILM Wako) on ice for 60 min. Cell lysates were then centrifuged at 12,000×*g* at 4°C for 20 min, diluted with ×4 SDS sample buffer (240 mM TrisHCl, pH 6.8, 8% SDS, 40% glycerol, 0.1% blue bromophenol, and 20% 2-mercaptoethanol), and heat-denatured at 97°C for 10 min. A cell lysate equivalent to 5×10^5^ cells/lane was loaded onto a 10% SDS polyacrylamide gel (ePAGEL E-T10, Atto). After electrophoresis, proteins were blotted onto Immobilon-P (Millipore) with a semi-dry blotting apparatus (TRANS-BLOT SD, Bio Rad). The membrane was blocked with TBS-T (50 mM TrisHCl pH 7.5, 150 mM NaCl, and 0.05% (v/v) Tween-20) containing 5% BSA at 4°C for 18 h. The membrane was incubated with the primary antibody in TBS-T containing 5% BSA (PY99: 1/1,000, LAT: 1/1,000 and β-actin: 1/2,000 dilution) at room temperature for 60 min, then washed with TBS-T three times. The membrane was incubated with the secondary antibody (HRP-conjugated anti-mouse IgG, diluted 1/3,000 in TBS-T containing 5% BSA) for 30 min and washed with TBS-T four times. Proteins and tyrosine-phosphorylated proteins were developed with SuperSignal Pico PLUS (Thermo Fisher) and visualized with a chemiluminometer (LuminoGraph 1, Atto).

### Flow cytometry

Cells were stained with the antibodies listed in the antibody table. 7-AAD or Zombie Violet (Biolegend) was used to discriminate between live/dead cells. To stain the transcription factors PLZF and RORγt, cells stained with the åsurface markers were fixed and permeabilized with the Transcription factor buffer set (BD Biosciences), and then stained with transcription factor antibodies. In CD107a staining, a fluorochrome-labeled CD107a antibody was added to the culture prior to the assay. Cells were analyzed with the MACSQuant cell analyzer (3 lasers, 10 parameters, Miltenyi Biotech) or the AttuneNxT acoustic focusing cytometer (4 lasers, 14 parameters, Thermo-Fisher Scientific). Data were processed using FlowJo software (ver.9.9 or 10.7, BD Biosciences). Cell sorting was performed using a FACSJazz cell sorter (2 lasers, 8 parameters, BD Biosciences).

### Cytokine and chemokine quantification

The quantification of cytokines and chemokines was performed with the LegendPlex mouse Th cytokine panel, mouse cytokine panel 2, and mouse proinflammatory chemokine panel according to the protocol provided by the manufacturer (Biolegend).

### Migration of m-reMAIT cells in syngeneic C57BL/6

Ly5.2-m-reMAIT cells (L7-1: 1.0 × 10^7^ cells/mouse) were intraperitoneally (i.p.) injected into C57BL/6 (Ly5.1) recipients and the expression of markers in endogenous (Ly5.1) and exogenous MAIT cells (Ly5.2 m-reMAIT cells) was assessed using a combination of appropriate antibodies and mMR1-tet 7 and/or 14 days after their transfer.

### Time course of MR1 expression in cancer cell lines upon the 5-OP-RU challenge

B16F10, EL4, LLC, RL♂1, and Yac-1 were incubated with 600 nM of 5-OP-RU. Thereafter, the geometric mean fluorescent intensity (gMFI) of MR1 on the cell surface was followed with by flow cytometry. gMFI at the indicated time point relative to that at the non-treated state (time 0) was calculated and shown as relative gMFI.

### Tumor studies

LLC suspended in HBSS was intravenously (i.v.) inoculated (3.0 × 10^5^ cells/mouse) 6 days after the adoptive transfer of m-reMAIT cells (1.0 × 10^6^ cells/mouse unless otherwise indicated, i.p.) or HBSS alone. In the survival assay, mice were considered to be dead when they showed a humane end-point, such as acute weight loss, hypothermia, and severe gait and/or consciousness disturbance, and the survival time of mice was plotted as a Kaplan-Meier curve. In the *in situ* tumor growth analysis, LLC (3.0 × 10^5^ cells/mouse) were subcutaneously inoculated 6 - 8 days after the adoptive transfer of m-reMAIT cells (0.3 × 10^6^, 1.0 × 10^6^, and 3.0 × 10^6^ cells/mouse, i.p.) or HBSS alone. Tumor sizes were measured with calipers every three days. Tumor volumes were calculated using the following formula: v= (tumor width)^2^ × (tumor length)/2.

### Interaction between NK cells and m-reMAIT cells

NK cells were isolated from C57BL/6 mice (male and female) with the MojoSort mouse NK cell isolation kit (BioLegend) (purity >85 % based on the flow cytometric analysis with NK1.1 and CD49b antibodies), and cocultured with m-reMAIT cells at 37°C in 5% CO_2_ for 18 h at the indicated ratio. In parallel, m-reMAIT cells and NK cells were cultured individually under the same conditions, and CD69 expression in NK cells (gated as NK1.1^+^CD49b^+^ cells) and m-reMAIT cells (gated as TCRβ^+^mMR1-tet^+^ cells) was analyzed using flow cytometry. Cytokines and chemokines released in the culture were measured with LegendPlex at the indicated ratio. In CD69 and CD107a expression analyses, the above column-isolated NK cells (1 × 10^5^ cells) and m-reMAIT cells (1 × 10^5^ cells, purity> 95%) were cultured individually or cocultured in the absence or presence of the same number of Yac-1 at 37°C in 5% CO_2_ for 18 h. The percentages of CD69^+^ cells and CD69^+^CD107a^+^ cells among NK cells and m-reMAIT cells were then measured using the MACSQuant flow cytometer.

### Semi-quantitative PCR

NK cells cocultured with m-reMAIT cells (NK cells/m-reMAIT cells ratio =1) were sort-purified as NK1.1^+^CD49b^+^ NK cells and TCR*β*^+^mMR1-tet^+^ m-reMAIT cells, respectively. RNA from each subpopulation was extracted with the RNeasy Mini kit (Qiagen), and cDNA was synthesized with the first-strand cDNA synthesis kit (Thermo Fisher Scientific) and then subjected to semi-quantitative PCR. PCR was performed with CYBR Green reagent (Nippon Genetics) using the following program: at 95°C for 5 min (90°C for 15 sec, 60°C for 60 sec) × 50 cycles (Light/Cycler Nano, Roche). The primer sets used in this study were described in the supplemental Table 1.

### Cytolytic activity

The cytolytic activities of NK cells and m-reMAIT cells were measured with Yac-1 and LLC as target cells. Yac-1 cells (1.0 × 10^4^ cells/assay) were labeled with CFSE 24 h prior to the addition of effector cells, while LLC cells (1.0 × 10^4^ cells/assay) were not labeled. Target cells were then incubated with the effector cells at the indicated E/T ratio at 37°C in 5% CO_2_ for 4 h and 16 h for Yac-1 cells and for LLC cells, respectively. E/T ratios of 1/1, 1/3, and 1/10 in the assay using the combination of NK cells and m-reMAIT cells contained twice the number of effector cells than NK cells or m-reMAIT cells alone. Dying target cells were stained with the Zombie Flexible Viability kit according to the manufacturer’s instructions. Percent lysis was calculated by the formula {(% of Zombie^+^ cells among target cells in the presence of effector cells) - (% of Zombie^+^ cells among target cells in the absence of effector cells)}/ 100 − (% of Zombie^+^ cells among target cells in the absence of effector cells)/100.

### Quantification and statistical analysis

Statistical analyses were conducted using Prism 9 for macOS (GraphPad). The Log-rank test was used for survival analyses between the two indicated groups. A two-way ANOVA was employed to assess the significance of differences among the various effector cells (NK cell, m-reMAIT cells, and NK cells plus m-reMAIT cells) in lysis assays.

### Data deposition

**Sequecnign data for TRBV of MAIT-iPSCs (Table 1) were deposited in DDJB under accession code**

LC637403 for L3-1 clone

LC637404 for L7-1 clone

LC637405 for L11-1 clone

LC637406 for L15-1 clone

LC637407 for L19-1 clone

## Acknowledgments

We thank M. Ohyama for technical help, Dr. Y. Horihata (Department of Biochemistry) for the structure analysis of synthesized 5-OP-RU, Y. Machida and T. Tsukada (Animal facility) for tumor experiments, Drs. S. Nishioka and M. Ikawa (Biken, Osaka University, Osaka, Japan) for chimeric mice, Drs. M. Nakanishi and M. Ohtaka (Tokiwa Bio. Co., Ltd., Tsukuba, Japan) for KOSM302L, Dr. X. Wang (Department of Pathology and Immunology, Washington University, MO, USA) for WT3, WT3/mMR1, CH27, and CH27/mMR1, Dr. T. Seya (School of Medicine, Hokkaido University, Sapporo, Japan) for B16F10, the NIH Tetramer core facility (Emory University, GA, USA) for unlabeled and APC- and BV421-labeled mMR1-tetramers, and Advanced Animal Model Support for supporting chimeric mice (JSPS KAKENHI 16H06276).

## Author Contributions

Conceptualization, H.W. and C. S.; Methodology, C. S., Y. M., H.W., and H.F.; Validation, C. S., H.W., and H.F.; Resources, C. S., H.W., and H.F.; Formal analysis, C.S.; Investigation, C.S., Y. M., H.W., and E.I.; Data curation, C.S., H.W., Y. M., and H.F.; Writing-original draft, H.W. and C.S.; Visualization, H.W. and C.S.; Supervision, H.W.; Project administration, H.W., H.F., and C.S.

Funding acquisition, H.W. The Science Research Promotion Fund 2018-2019 (The Promotion and Mutual Aid Corporation for Private Schools of Japan); C. S. 17H03565 (JSPS KAKENHI) and 26430084 (JSPS KAKENHI); H. F. 20K10435 (JSPS KAKENHI).

## Legends for Supplementary Figures

**Figure S1.**
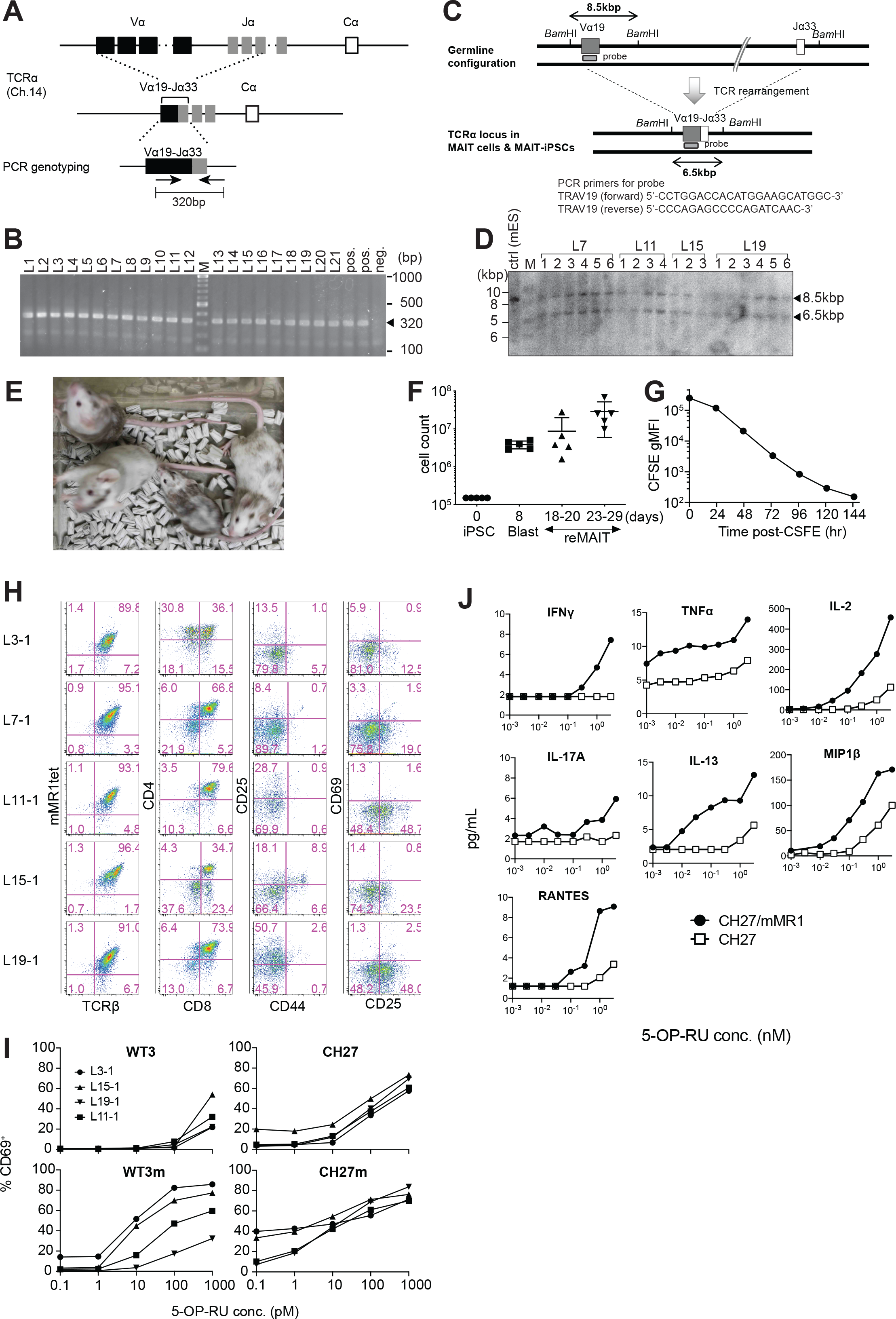
iPS cells (iPSCs) from MAIT cells and m-reMAIT cells (related to Figure 1. **S1A. Genomic configuration of *TCRα* loci in MAIT-iPSCs** *TCRα* loci in C57BL/6 (upper panel) and MAIT-iPSCs (middle panel). The primer set to detect rearranged TRAV1-TRAJ33 (Vα19-Jα33) is shown with arrows (lower panel). **S1B. MAIT-iPSC detection by PCR** The results of PCR using the primer set shown in A. Bands corresponding to rearranged *TRAV1-TRAJ33* (*Vα19-Jα33*) are indicated by arrowheads. MAIT-iPSCs (L1-L21) before the limiting dilution, the positive control, and negative control (ES cells) are shown. M, molecular weight marker **S1C. Southern blot probe detecting the rearranged *TCRα* locus** The genomic configuration of the *TCRα* locus before (upper panel) and after (lower panel) the Vα19-Jα33 rearrangement. The positions of the Southern blot probe as well as *BamH*I sites are shown together with the primer sequences for probe synthesis. **S1D. Southern blot analysis of MAIT-iPSCs** The bands corresponding to the germline configuration (∼8.5 kb) and those after the rearrangement (∼6.5 kb) are shown by arrowheads. In the analysis, genomic DNA from MAIT-iPSCs after limiting dilutions was used. Mouse ES cells (cntrl), M: DNA size marker, L7 clones; L7-1∼L7-6, L11 clones; L11-1∼L11-4, L15 clones; L15-1∼L15-3, and L19 clones; L19-1∼L19-6 **S1E. Chimeric mice generated from MAIT-iPSCs** Chimeric mice from MAIT-iPSCs (L7) are shown. Chimerism ranged between 10 and 90%. **S1F. Expansion of MAIT-iPSCs during differentiation** Cell numbers at the indicated differentiation stage of L7-1 from iPSCs are plotted. Blast: lymphocyte progenitor cells, reMAIT: m-reMAIT cells defined as TCRβ^+^mMR1-tet^+^ Data are shown as medians. Horizontal line: Median; whiskers: Minimum and Maximum. **S1G. m-reMAIT cell doubling time during the logarithmic phase of expansion** The fluorescent intensity of CFSE-labeled m-reMAIT cells (L7-1) vs. the culture time is plotted to estimate the doubling time. **S1H. Flow cytometric profiles of m-reMAIT cells from different MAIT-iPSCs** mMR1-tet staining and the expression of TCRβ, CD4, CD8, CD25, CD69 and CD44 in m-reMAIT cells (from L3-1, L7-1, L11-1, L15-1, and L19-1 MAIT-iPSCs) on differentiation day 23 **S1I. 5-OP-RU dose-dependent activation of m-reMAIT cells** The percentage of CD69^+^ cells among m-reMAIT cells (from L3-1, L11-1, L15-1, and L19-1 MAIT-iPSCs) challenged with the indicated concentration of 5-OP-RU in the presence of APCs, such as WT3, WT3/mMR1, CH27, and CH27/mMR1, is shown. **S1J. 5-OP-RU dose-dependent production of cytokines and chemokines** The indicated cytokines and chemokines in the culture supernatant from m-reMAIT cells (L7-1) cultured in the presence of CH27 (□) or CH27/mMR1 (●), and CHALLENGED with varying concentrations of 5-OP-RU are quantified with LegendPlex. Data are representative of two independent experiments.

## Legends for Supplementary Tables

**Supplemetary Table 1.**
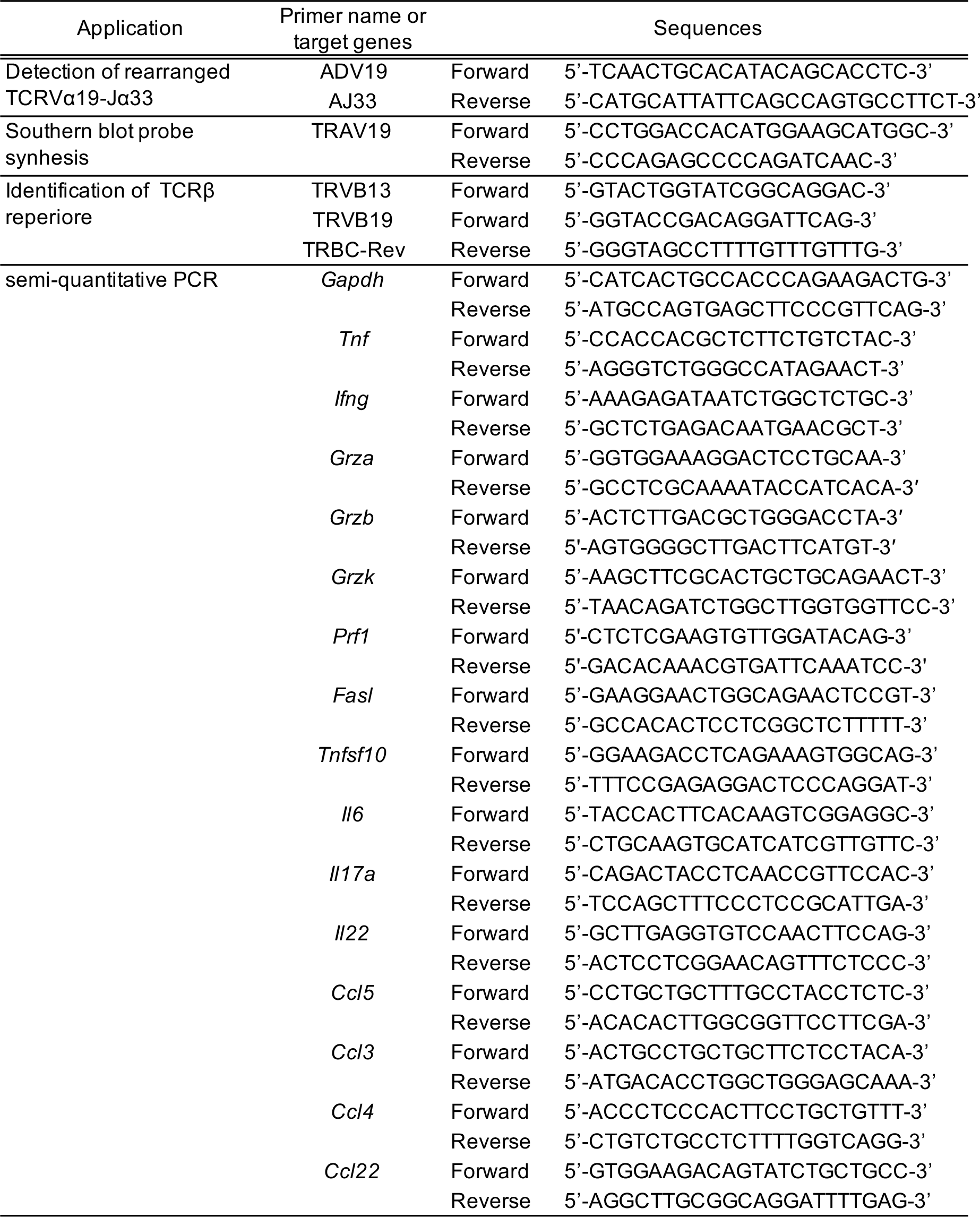
Oligonuleotides used in this study.

**Supplemetary Table 2.** Antibodeis used in this study.

| Application | Antibodies | Vendors |
| --- | --- | --- |
| Flow cytometry | CD4 (RM4-4), PerCP/Cy5.5 | BioLegend |
| | CD8α (53-6.7), APC/Cy7 | BioLegend |
|  | CD25 (PC61), APC | BD Biosciences |
|  | CD25 (PC61), APC | BioLegend |
|  | CD25 (PC61), BV421 | BioLegend |
|  | CD25 (PC61), BV711 | BD Biosciences |
|  | CD25 (PC61), PE | BioLegend |
|  | CD44 (IM7), FITC | BioLegend |
|  | CD45.1 (Ly5.1) (A20), BV605 | BioLegend |
|  | CD45.2 (Ly5.2) (104), PE | BioLegend |
|  | CD45R/B220 (RA3-6B2), BV421 | BioLegend |
|  | CD45R/B220 (RA3-6B2), PE | BioLegend |
|  | CD49b (DX5), PE | BioLegend |
|  | CD69 (H1.2F3), APC/Cy7 | BioLegend |
|  | CD69 (H1.2F3), PE | BD Biosciences |
|  | CD69 (H1.2F3), PE/Cy7 | BioLegend |
|  | CD107a (1D4B), BV421 | BioLegend |
| | CD127 (IL-7Rα) (A7R34), PerCP/Cy5.5 | BioLegend |
|  | CD186 (CXCR6) (SAO51D1), PE | BioLegend |
| | CD218a (IL-18Rα) (P3TUNYA), eFluor450 | Thermo Fisher Scientific |
| | CD218a (IL-18Rα) (REA947), PE | Miltenyi Biotech |
|  | F4/80 (BM8), BV421 | BioLegend |
|  | F4/80 (BM8), PE | BioLegend |
|  | MR1 (26.5), APC | BioLegend |
|  | MR1 (26.5), purified | BioLegend |
|  | NK1.1 (PK136), BV510 | BD Biosciences |
|  | NK1.1 (PK136), FITC | BioLegend |
|  | PLZF (9E12), PE | BioLegend |
| | RORγt (Q31-378), BV421 | BD Biosciences |
| | TCR Vβ6 (RR4-7), PE | BioLegend |
| | TCR Vβ8.1, 8.2 (MR5-2), PE | BioLegend |
| | TCRβ (H57-597), BV605 | BD Biosciences |
| | TCRβ (H57-597), BV605 | BioLegend |
| | TCRβ (H57-597), PE/Cy7 | BioLegend |
|  | American Hamster IgG isotype control (HTK888), PE | BioLegend |
|  | Mouse IgG2a, k isotype control (MOPC-173), BV421 | BioLegend |
|  | Mouse IgG2a, k isotype control (MOPC-173), purified | BioLegend |
| Western blotting | β-actin (2F1-1), purified | BioLegend |
|  | LAT (11B.12), purified (for WB) | Santa Cruz |
|  | Phosphotyrosine (PY99), purified (for WB) | Santa Cruz |
| in vivo depletion | Mouse, Rat Asialo GM1 | FUJIFILM Wako |

